# Single cell landscape of sex differences in the different courses of multiple sclerosis

**DOI:** 10.1101/2024.06.15.599139

**Authors:** Irene Soler-Sáez, Borja Gómez-Cabañes, Rubén Grillo-Risco, Cristina Galiana-Roselló, Lucas Barea-Moya, Héctor Carceller, María de la Iglesia-Vayá, Sara Gil-Perotin, Vanja Tepavčević, Marta R. Hidalgo, Francisco García-García

## Abstract

**Background:** One of the major challenges in addressing multiple sclerosis is to understand its progression trajectory. The pathological process transitions from acute phases predominantly driven by inflammation to progressive clinical profiles where neurodegeneration takes precedence. It is known that sex plays a crucial role in this heterogeneity; females are two to three times more likely to suffer from multiple sclerosis, while males suffer from more rapid neurodegeneration with greater severity.

**Results:** To gain insight into the sex-based molecular differences, we processed single cell datasets from the central nervous system and the peripheral blood, covering the different courses of multiple sclerosis. We generated cell-type specific landscapes, including gene signatures from differentially expressed genes, functional profiling, pathway activation, and cell-cell communication networks for females, males, and their sex differential profiles. Among our findings, we revealed that female neurons may exhibit protective mechanisms against neurodegeneration. In the inflammatory-predominant forms, female immune cells present an inflammatory core driven by the AP-1 transcription factor, while male adaptive immune cells exhibit higher mitochondrial impairment. Conversely, larger differences are reported in CD8+ T cells progressive forms, with males exhibiting cytolytic profiles that may promote neurodegeneration. Complete results can be explored in the interactive webtool https://bioinfo.cipf.es/cbl-atlas-ms/.

**Conclusions:** We identified cell-type specific sex differences in brain and immune cells that vary in the spectrum of multiple sclerosis. We consider this molecular description a valuable resource to promote future targeted approaches considering the sex of the individual.

## BACKGROUND

Multiple sclerosis (MS) is a chronic, autoimmune, and neurodegenerative condition of the central nervous system (CNS) that represents the leading cause of non-traumatic neurological disability in young adults (25-30 years).^1–3^ MS disability course represents a continuum of molecular and cellular events that accentuate and compensate for the disease’s clinical traits. These events are driven by interconnected neurodegenerative, inflammatory, and reparative mechanisms.^4^ Defective autoimmune responses partly determine myelin destruction and axonal damage. Moreover, prolonged deviation from CNS physiological norms also promotes hyperexcitability and reactive oxygen species generation, which enhances neurodegeneration. Astrocytes, microglia, and oligodendrocytes play dual roles involving both protective (e.g., myelin recovery and maintenance of lipid homeostasis) and harmful (e.g., enhanced proinflammatory environment) activities.^5,6^

MS progression has been classified for ease of communication. The most common form - relapsing-remitting MS (RRMS) - fluctuates between periods of relapses, driven by exacerbated inflammatory status and followed by subsequent remissions. Evolution into a progressive course gives rise to secondary progressive MS (SPMS). Lastly, primary progressive MS (PPMS) is characterized by a chronic neurological decline from the onset, predominantly driven by neurodegeneration and not preceded by relapses.^7^

Sex contributes as an important variable to epidemiological and clinical aspects. Females tend to develop MS earlier, exhibit a higher prevalence (with a ratio of 2-3:1), and suffer more severe inflammatory patterns.^8^ In contrast, males experience more rapid CNS deterioration, with more significant atrophy.^9^ These differences are reflected in the clinical categorization of the disease. Females are more prone to suffer RRMS (with more relapses), while the progressive forms appear at a more balanced ratio between the sexes.^10^ Studies in human and model organisms have sought to unravel these sex-driven disparities, primarily focusing on specific genes or molecular mechanisms.^11^

We aimed to uncover novel insight into how sex differences manifest in MS, providing a more profound knowledge of the mechanisms driving such disparities. We report comprehensive atlases of sex differences in MS subtypes by analyzing single-cell RNA sequencing (scRNA-seq) and single-nucleus RNA sequencing (snRNA-seq) datasets from peripheral blood mononuclear cells (PBMCs) and CNS samples, respectively. Neurons, astrocytes, microglia, oligodendrocytes and oligodendrocyte precursor cells (OPCs) were identified for CNS samples; and CD4+ T cells, CD8+ T cells, NK cells and monocytes for PBMCs. We examined sex-specific and sex differential changes in gene expression patterns, functional profiles, signaling pathways, and cell-cell communication. Our results reveal significant transcriptomic alterations that could explain sex differences in MS progression, such as potential compensatory mechanisms to cope with excitotoxicity in female neurons and higher activation of male CD8+ T cells during disease progression. These findings may have important implications to better understand the disease, taking them into account when designing future strategies to enhance neuroprotection, control inflammation, promote myelin recovery, and delay neurodegeneration.

## RESULTS

### All cell types exhibit transcriptomic sex differences across the course of multiple sclerosis

We analyzed three different datasets to obtain the sex-differential landscape of MS subtypes. Table S1 summarizes the characteristics of each dataset, including the number of samples per group and the number of cells and genes evaluated. The systematic review results are reported in Figure S1 and the Supplementary Note includes an extended description of all datasets analyzed in this work. We processed all datasets independently, with results shown in Figure S2-S4. Data processing and cellular annotation steps led to the identification of the major cell types in each tissue. For each cell type, we identified those genes with different expression patterns (differential gene expression analysis), in which functions are these genes involved (functional profiling analysis), and the differential activation of signaling pathway effectors (signaling pathways analysis). We evaluated all biological inference approaches in three different scenarios: impact of disease in females (IDF: MS.Female - Control.Female), impact of disease in males (IDM: MS.Male - Control.Male), and sex-differential impact of disease (SDID: (MS.Female - Control.Female) - (MS.Male - Control.Male)). We also performed cell-cell communication analyses to quantitatively infer interaction networks for each group (MS.female, Control.Female, MS.Male, and Control.Male). As a result, we found significant differences in all comparisons and biological inference approaches, reflecting the impact that sex may have on the course of the disease. We elaborate the results in the following sections and comprehensively report them in https://bioinfo.cipf.es/cbl-atlas-ms/.

### Atlas of sex differences in secondary progressive MS *post-mortem* brain tissue

We identified neurons, astrocytes, microglia, oligodendrocytes, and OPCs within brain samples (Figure 1A). Figure S5A describes the expression pattern of the marker genes, while Figure S5B describes the number of cells by type, condition, and sex. We encountered sex differences in all cell types (Figure 1B). Although many genes, functions, and pathways remained specific to one sex, a considerable number of them presented simultaneous alterations in both sexes, creating a catalog of different patterns (Figure 1C). We focused on the features significant in all three comparisons (Figure 1C, colored combinations), that is, features that significantly change in the disease in females (IDF comparison) and males (IDM comparison), but also present significant sex differences in the SDID comparison. These alterations fall into six patterns, which we then grouped into two by their logFC sign in the SDID comparison: i) significant features increased in females (logFC > 0), and ii) significant features increased in males (logFC < 0) (Figure 1D). Henceforth, we used this classification to illustrate results for simplicity’s sake. Figure S6-10 and Table S2 summarize significant features for each cell type, including volcano plots for each comparison, PPI networks and semantic clustering of significant functions. Most features display cell type-specificity or, at most, were significant in two different cell types (Figure S11).

**Figure 1.**
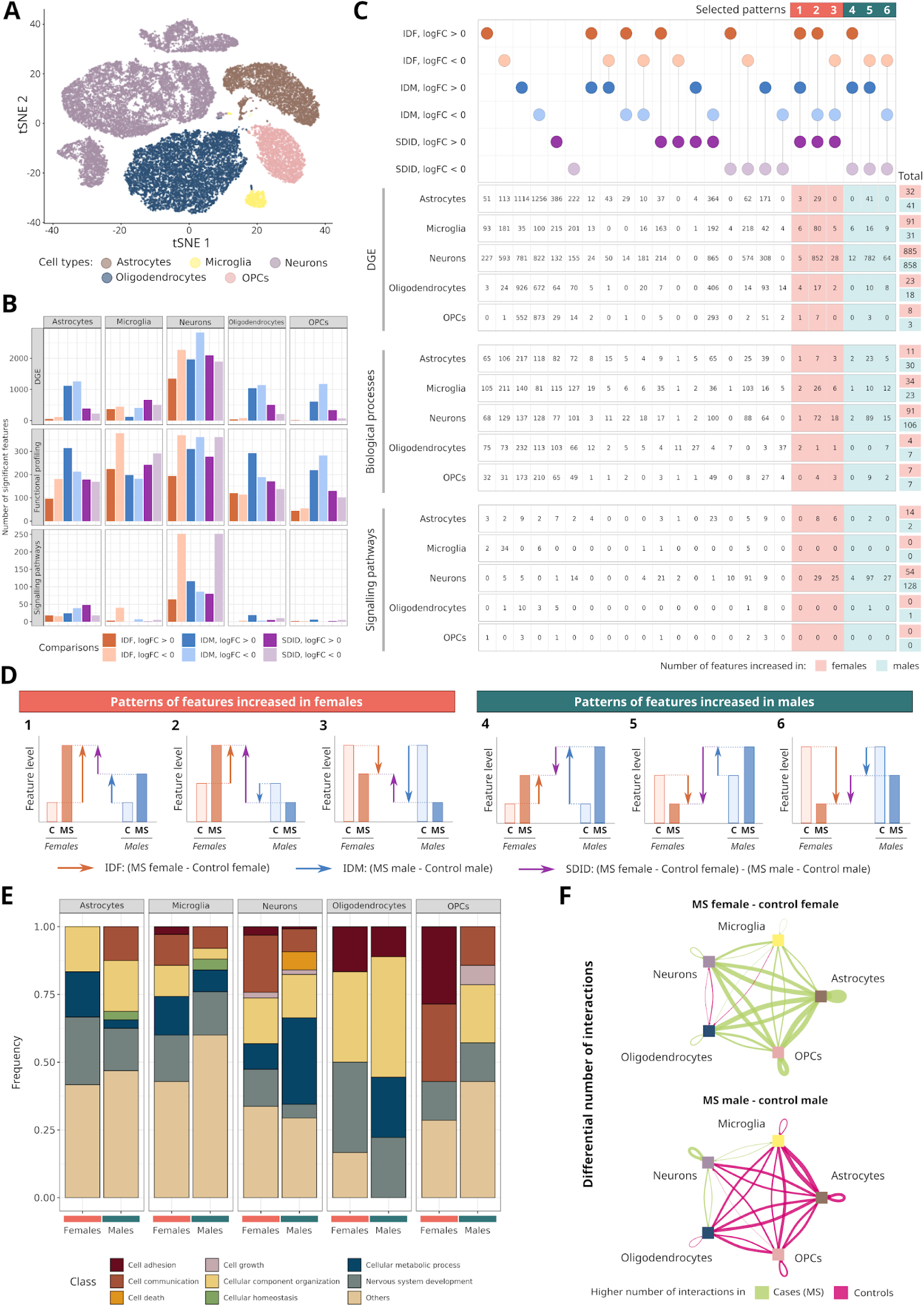
Transcriptomic landscape of sex differences in secondary progressive MS central nervous system. (**A**) Cell type distribution in tSNE dimensions. Each dot represents a cell, colored by annotated cell type. (**B**) The number of significant features by cell type, analysis, and comparison. For functional profiling, it represents the number significantly enriched BP-GO biological processes (**C**) Significant features for each cell type separated by comparison and direction of change (logFC). Dots connected by vertical lines denote comparisons in which the feature is statistically significant, with specific colors indicating the comparison and logFC sign. Each row of numbers indicates the number of significant features corresponding to the evaluated column in the dot map for each cell type and specific analysis. Colored squares highlight significant features in the IDF, IDM, and SDID comparisons, corresponding to (**D**) six qualitatively detailed patterns. (**E**) The relative frequency distribution of BP-GO terms significantly overrepresented in females (orange) and males (green) by cell type. Individual terms classified into general categories (See legend). (**F**) Differential cell-cell communication networks between MS females and control females (top) and MS males and control males (bottom). Colors indicate more interactions in MS (green) or controls (pink); the thickness of the interaction corresponds to the magnitude of change. *BP-GO: biological processes from Gene Ontology; C: controls; DGE: differential gene expression; IDF: impact of disease in females; IDM: impact of disease in males; MS: multiple sclerosis; OPCs: oligodendrocyte precursor cells; SDID: sex differential impact of disease; tSNE: t-Distributed Stochastic Neighbour Embedding*.

In general terms, females displayed a higher proportion of functions related to *cell adhesion* (microglia, neurons, oligodendrocytes, and OPCs) and *nervous system development* (astrocytes, neurons, and oligodendrocytes) (Figure 1E). Conversely, males exhibited a higher proportion of functions implicated in *cell death* (neurons), *cellular homeostasis* (astrocytes and microglia), and *cell growth* (OPCs) compared to females. Concerning significant pathway effectors, the most frequent terms increased in females related to *Infection diseases* and *Cell growth and death* (astrocytes) and *Signal transduction* (neurons), while male effectors mainly related to *Signal transduction* (astrocytes) and *Nervous system* (neurons) (Figure S12A).

Cell-cell communication analyses complete our atlas of sex differences. Figure S12B describes the number of inferred interactions. Notably, males and females displayed practically opposite disease patterns (Figure 1F): females presented a greater number of cell-cell interactions in MS, while males in healthy controls. Remarkably, we observed increased neuron-neuron interactions in MS males and MS females, although the value for males exhibits a larger magnitude of change.

#### Sex differential alterations in the astrocyte-microglia-neuron triad of secondary progressive MS implicate synaptic components and stress responses

We performed a semantic clustering analysis for each sex over the enriched biological functions in astrocytes, neurons, and microglia. This approach allowed us to determine whether similar functions were enriched in different cell types or, on the contrary, if most of them were cell type-specific functional changes. Results showed that functional alterations in both males and females were distributed across similar semantic areas, without a clear distinction in clustering patterns pointing to a specific cell type (Figure 2A). Notably, the comparison of the word cloud between males and females revealed certain functional similarities. Biological functions related to RNA processing were found to be significant in both sexes (female cluster 4 and male cluster 1), although males specifically presented enriched functions related to protein degradation and apoptosis (male clusters 1 and 3). Table S3 and Table S4 provide detailed descriptions of these functions.

**Figure 2.**
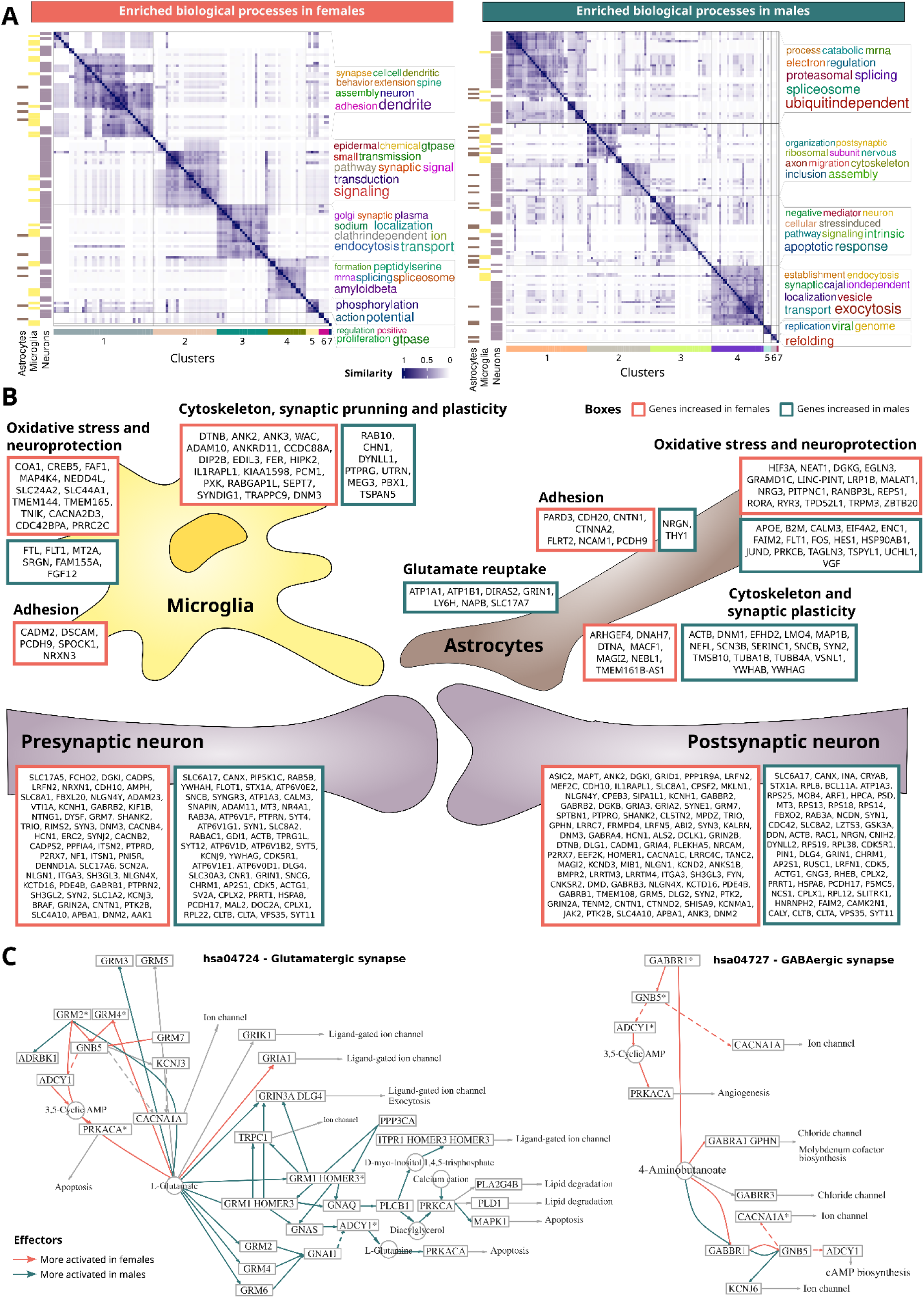
Sex differences in secondary progressive MS post-mortem brain tissue synapses. (**A**) The degree of similarity between BP-GO terms overrepresented in females (left) and males (right) for astrocytes, microglia, and neurons. Each row and column corresponds to a significant BP-GO term. The intensity of blue shading reflects the semantic similarity between terms. The horizontal bars on the left indicate whether the term is significant in astrocytes, microglia, and/or neurons. The clusters visualized at the bottom correspond to the word cloud displayed to the right of the plot. (**B**) Atlas of sex differences in synapse-related gene expression. Significant genes involved directly/indirectly in synapses were selected and classified in broad categories based on their roles in the cells. Colored boxes indicate sex patterns: genes upregulated in females (orange) and males (green). (**C**) Signaling pathways of glutamatergic (left) and GABAergic (right) synapses. Nodes represent proteins of the signaling pathway and edges the interactions between nodes. Effector proteins are the last node in each subpathway (arrow point). The effector nodes point to the biological functions they exert. *BP-GO: Biological processes from Gene Ontology; GABA: Gamma-aminobutyric acid*.

The remaining clusters, both in female- and male-enriched scenarios, were predominantly associated with synaptic processes, suggesting that sex-related differences may affect the synaptic transmission. To further characterize these differences, Figure 2B summarizes the differentially expressed genes associated directly/indirectly with synaptic features. Neurons possess a large number of presynaptic and postsynaptic terms with sex differential dysregulation; some gene families display a consistent increase in the same sex, such as voltage-gated calcium channels in females (CACNA1C, CACNB2, and CACNB4) and ATPase subunits involved in maintaining ion gradients, vesicular acidification, and neurotransmitter packaging in males (ATP6V0E2, ATP1A3, ATP6V1F, ATP6V1G1, ATP6V1D, ATP6V1B2, ATP6V1E1, and ATP6V0D1). Female neurons also increased higher number of genes to modulate excitability such as glutamate (e.g., GRIA2 and GRIA3), and GABA receptors (e.g., GABRB1 and GABBR2). Accordingly, most glutamatergic effectors (increase excitability) display greater activation in males (Figure 2C, left); meanwhile, most GABAergic effectors (decrease excitability) display greater activation in females (Figure 2C, right). Importantly, we have validated these pathways in an independent cohort published by MacNair *et al.*,^12^ which revealed that 79% glutamatergic effectors and 86% of GABAergic effectors showed the same significant sex-differential activation pattern (Figure S13). Cholinergic, serotonergic, and dopaminergic synapses present further significant differences (Figure S14), sparking an intricate network of sex disparities in neuronal excitability.

Astrocytes and microglia presented significant differences in gene expression profiles associated with neuroinflammation and tissue homeostasis (Figure 2B, Oxidative stress and neuroprotection and Glutamate reuptake boxes). Intriguingly, male astrocytes express increased levels of glutamate reuptake-related genes. Sex differential stress responses in astrocytes also included the increased expression of genes related to calcium homeostasis (RYR3, TPD52L1, and TRPM3 in females and CALM3 in males), oxidative stress (REPS1 in females; ENC1, HSP90AB1, PRKCB, and UCHL1 in males), hypoxia (HIF3A, EGLN3, and ZBTBW in females; EIF4A2 in males) and neuroprotection (NRG3, RORA, NEAT1, RANBP3L, MALAT1, and LINC-PINT in females; JUND, FOS, FLT1, FAIM2 and HES1 in males). Meanwhile, female microglia notably increased the expression of genes associated with transmembrane transporters involved in metabolism and homeostasis maintenance, such as SLC24A2 (calcium/cation antiporter), SLC44A1 (choline transporter), TMEM144 (carbohydrate transporter), TMEM165 (cation/proton antiporter) and CACNA2D3 (calcium transporter), while male microglia increased the expression of genes associated with intracellular metal homeostasis (FTL and MT2A). Additionally, female and male microglia displayed increased expression levels of oxidative stress-related genes (COA1, CREB5, and FAF1 in females; SRGN and FGF12 in males). Glial cells also exhibited the dysregulated expression of genes potentially involved in myelin recovery, which are illustrated in the following section.

#### Sex differential alterations in secondary progressive MS post-mortem brain tissue also affect lipid metabolism and myelin recovery

We next explored oligodendrocytes and OPCs to gain insight into the sex differential potential for myelin repair and its maintenance. Neuronal-related functions and myelination represented the major altered categories in both cell types (Figure S15); however, associated genes differ between sexes (Table S5). Oligodendrocytes differed in genes encoding neuronal adhesion and synaptic molecules that support the maintenance of myelin targeting to axons, with NCAM2, NLGN1, CADM2, CLDN11, ANK3, IL1RAPL1, CTNND2, and LPHN3 increased in females and CAMK2B and NRGN in males. Females also displayed the increased expression of genes boosting myelin repair, such as AMER2 (negative regulator of the Wnt/ß-catenin pathway) and OMALINC (lncRNA marker of oligodendrocyte maturation that regulates myelination-associated gene expression). Interestingly, males also displayed the increased expression of marker genes for oligodendrocyte differentiation (CAMK2B and MIR219A2) and myelinating activity (MAPB1, BCAS1, and MGAT5).

Among the genes upregulated in female OPCs, we encountered genes involved in differentiation to oligodendrocytes and myelin repair processes (GPNMB, PARD3, QKI, and TNR) and neuroprotection (HIF3A, and VCAN), while male OPCs only display one gene (MAP1B) related with these functions. We next characterized PDGF (Platelet-derived Growth Factor) and FGF (Fibroblast growth factor) communication signaling to OPCs (Figure 3), given that these two pathways drive OPCs to growth and differentiate into oligodendrocytes to promote myelin repair. The PDGF and FGF signaling displayed higher interaction strengths in MS males and MS females compared to corresponding controls (Figure 3A-B, left). However, the contribution of donor cells and the strength of ligand-receptor pairs differed according to sex, with more pronounced alterations observed in MS females (Figure 3A-B, right). In PDGF signaling, the MS female PDGFC (OPCs) - PDGFRA (OPCs) interaction increased proportionally compared to control females to the detriment of interactions with neurons-OPCs. In FGF signaling, ligands provided by oligodendrocytes increased in interaction strength proportionally in MS females compared to female controls, while astrocyte ligand interactions decreased. Notably, we did not observe these patterns in PDGF and FGF signaling when comparing MS males to male controls.

**Figure 3.**
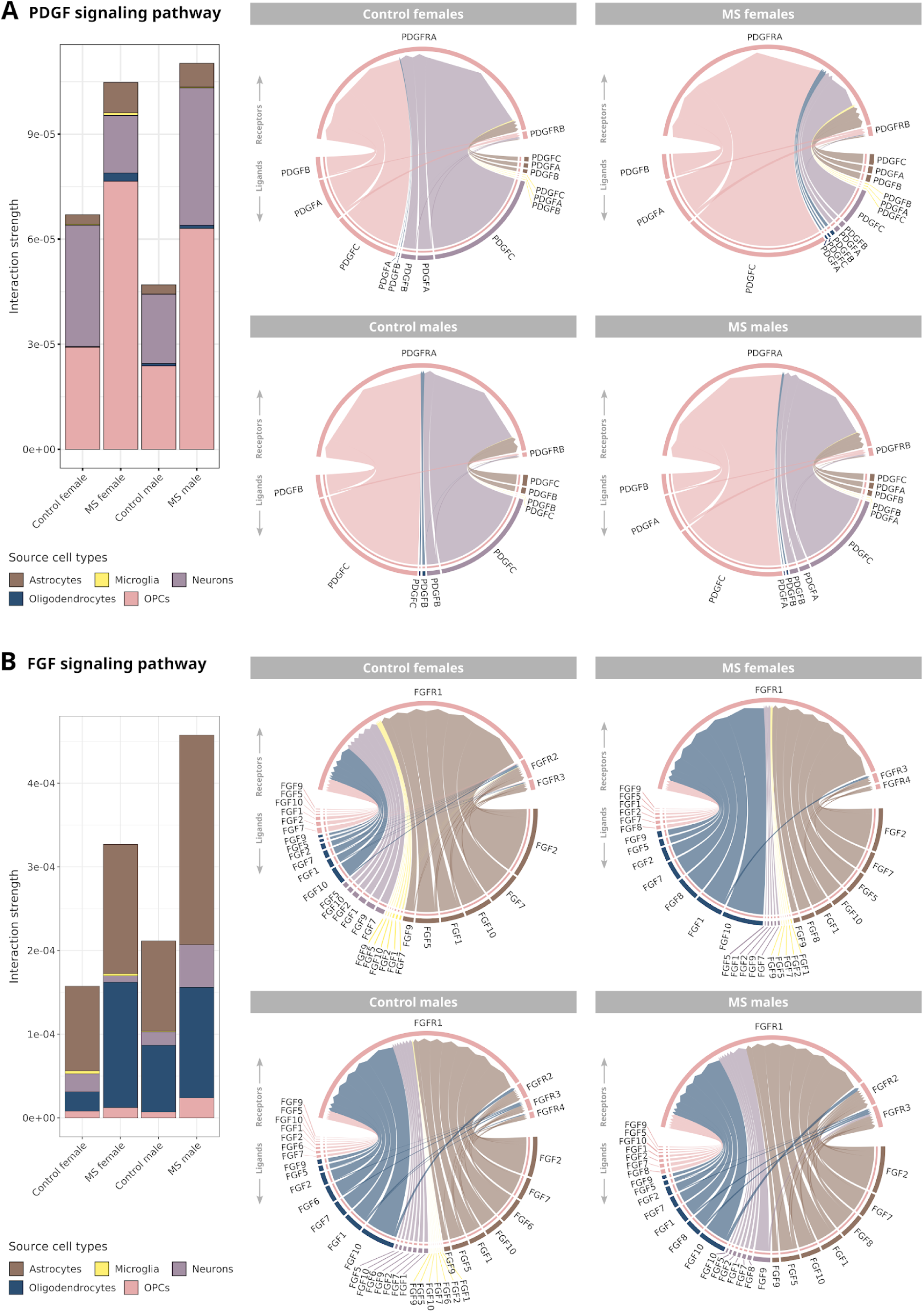
(A) PDGF and (B) FGF signaling received by oligodendrocyte precursor cells to promote their growth and differentiation. (Left) Total interaction strength by cell type for each interest group: MS females, control females, MS males, control males. (Right) Interaction strength frequency of ligand-receptors pairs. The outer circumference labels the cell type. The inner flux represents the proportional interaction strength colored by the sender cell type, which provides the ligand. Values represent the interaction strength inferred by CellChat algorithm, calculated based on the probability of communication between ligand–receptor pairs. Higher interaction probabilities correspond to greater signaling intensity. *Sender cells: astrocytes, microglia, neurons, oligodendrocytes, and oligodendrocyte precursor cells (OPCs). Receiver cells: OPCs. FGF: fibroblast growth factor; MS: multiple sclerosis; OPCs: oligodendrocyte precursor cells; PDGF: platelet-derived growth factor*.

Astrocytes and microglia also displayed significant sex differences in the expression of lipid metabolism-associated genes, which may promote myelin clearance. We observed more genes with increased expression in female than male astrocytes (e.g., DGKG, GRAMD1C, LRP1B, and PITPNC1). However, males displayed APOE increased expression, which could potentiate lipid droplet accumulation. Meanwhile, female microglia displayed the increased expression of genes such as CELF2, CHST11, AGO3, ATG4C, GPM6B, LPAR1, LPGAT1, OSBPL1A, PIP4K2A, PLD1, QKI, SOX2-OT, and TMEM131, while male microglia increased the expression of RAB10, TNS3, COLEC12, PLSCR1, SCARB1, and VIM. These results suggest adaptive differences in lipid metabolism and debris clearance from damaged myelin in MS.

### Atlas of sex differences in relapsing-remitting MS peripheral blood mononuclear cells

Next, we focused on B cells, CD4+ T cells, CD8+ T cells, NK cells, dendritic cells and monocytes in RRMS (Figure 4A). Figure S16 contains marker gene expression data, cell distribution by group and sex, and the number of statistically significant features; Figures S17 to S21 include an overview of the results for each cell type. We illustrated significant features in our three comparisons as previously defined in Figure 1D. Due to the limited number of results obtained for B cells and dendritic cells, we focused on CD4+ T cells, CD8+ T, and NK cells and monocytes (Figure 4A, colored boxes). In all cell types, females exhibited a higher number of significant functions than males in mostly all categories (Figure 4B), which agrees with the higher number of genes with significantly increased expression compared to males (Figure 4C). We noted an exception in the *Metabolism and bioenergetics* category for male CD4+ T, CD8+ T, and NK cells, which relates to mitochondrial electron transport. Lastly, the differential activation of signaling pathways and cell-cell communication interactions comprised an intricate framework of sex differences (Figure S22).

**Figure 4.**
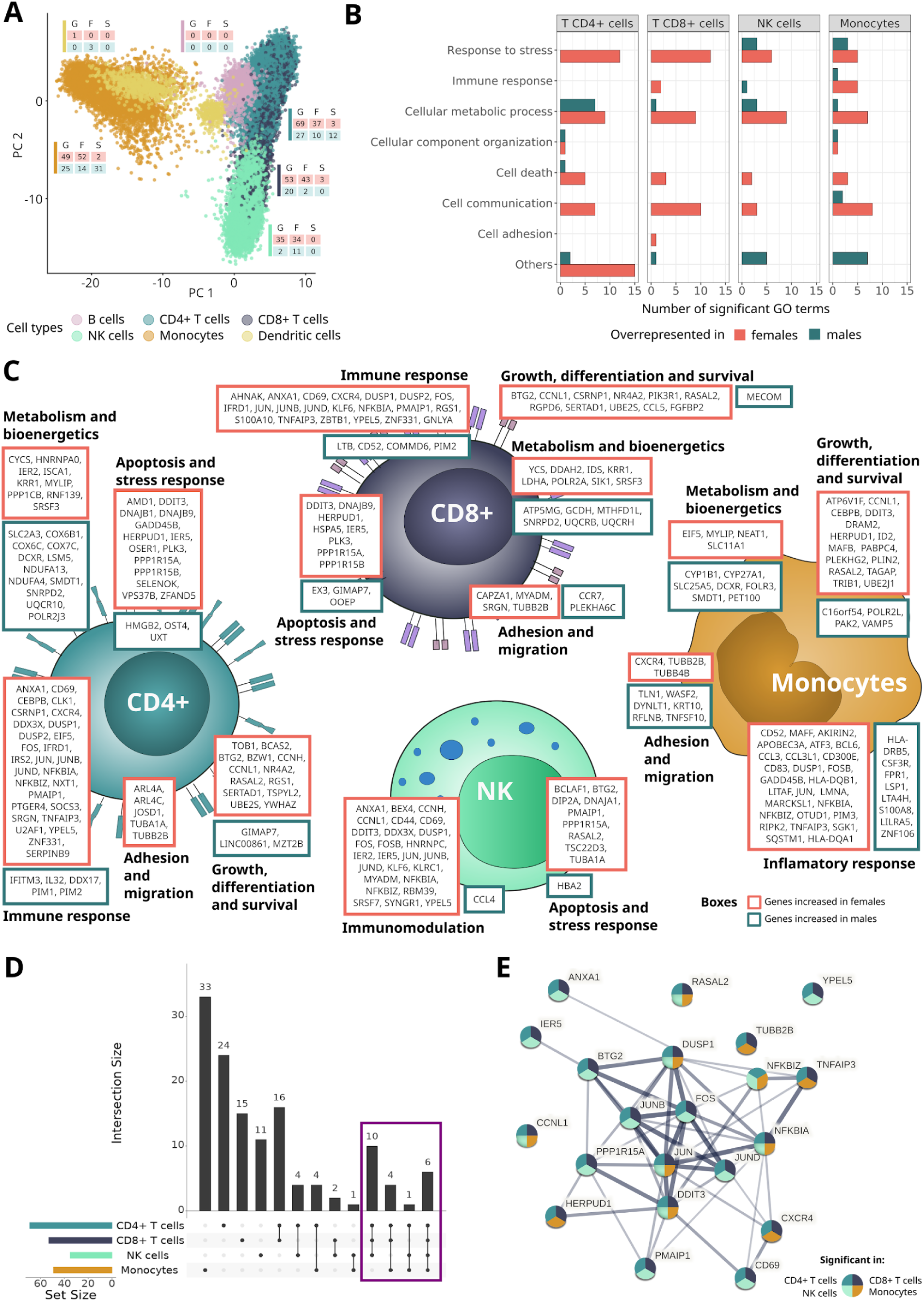
Transcriptomic landscape of sex differences in relapsing-remitting MS peripheral blood mononuclear cells. (**A**) Cell type distribution in PCA dimensions. Each dot represents a cell, colored by the annotated cell type. Colored boxes indicate that the number of genes (G), functions (F), and signaling pathways (S) significantly increased in females (orange) and males (blue). (**B**) The number of significant BP-GO terms by cell type classified under broader biological categories. (**C**) Atlas of sex differences in gene expression patterns. Genes classified in general biological terms. Colored boxes indicate the sex patterns: genes upregulated in females (orange) and males (green). (**D**) Upset plot of genes with increased expression in females compared to males by cell type. Horizontal bars represent the number of significant genes in each cell type. Dots indicate the combinations of intersections evaluated, with the top vertical bars denoting the number of significant genes of the corresponding intersection. Purple square: significant genes in at least three evaluated cell types. (**E**) Protein-protein interaction networks of significant genes with increased expression in females compared to males in at least three cell types evaluated: CD4+ T cells, CD8+ T cells, NK cells and monocytes. Edge thickness indicates the structural and functional confidence of the interaction. *BP-GO: Biological processes from Gene Ontology; NK: natural killer*.

#### Functional profiling reveals an immune signature core of relapsing-remitting MS females but not in MS males

The female peripheral immune system displays a greater number of genes with increased expression than males (Figure 4C). We discovered 21 significant genes in at least three of the four cell types evaluated (CD4+ T cells, CD8+ T cells, NK cells and monocytes), providing a general insight into the state of the female innate and adaptive immune systems (Figure 4D). These genes constitute a significantly connected network (p-value: < 1.0e-16) (Figure 4E), where the prominent hub comprises the AP-1 transcription factor complex (FOS, JUN, JUNB, and JUND). We did not identify a common gene core for males; the C16orf54 gene (associated with immune infiltration) represents the only significantly dysregulated gene in CD4+ T cells, CD8+ T cells and monocytes (Figure S23). Alterations in the transcriptomic profile were reflected in the differential interaction strength of ligand-receptors pairs in MS females compared to control females, where we detected the increased interaction of signaling pathways mediated by proinflammatory cytokines (e.g., IL6) and cell adhesion molecules (e.g., CD99 and cadherins) that do not become reinforced in males (Figure S22B).

#### The adaptive immune response in relapsing-remitting MS males exhibits exacerbated mitochondrial dynamics compared to females

We previously noted the absence of a central hub of genes with increased expression in the immune system of MS males; however, we found intriguing results by separating the innate and adaptive immune systems. The adaptive immune system presents different genes functionally related to the mitochondrial electron transport chain (Figure 4C). CD4+ T cells displayed the increased expression of genes from all steps: NADH to ubiquinone (NDUFA13 and NDUFA4), ubiquinol to cytochrome c (UQCR10), and cytochrome c to oxygen (COX6B1, COX6C, and COX7C), while CD8+ T cells displayed the increased expression of genes for two additional ubiquinol-cytochrome C reductases (UQCRB and UQCRH) and ATP5MG, a gene involved in ATP synthesis coupled with proton transport. These genes could be activating to a greater extent mitochondrial electron transport chain in mitochondria in males compared to females, with a potential increase in reactive oxygen species that may enhance neurodegeneration. Meanwhile, male monocytes display an elevated activation of pathways primarily involved in cell adhesion and inflammatory modulation compared to females (Figure S22A). Male monocytes also increase the expression of genes that favor shape-shifting infiltration (DYNLT1, TLN1, and RFLNB) and mediate innate inflammation (LTA4H, S100A8, and LILRA2) (Figure 4C). Notably, we observed an increase in the interaction of receptor-ligand pairs considering all cell types, which involved response modulation via semaphorins and the T-lymphocyte co-stimulatory molecules CD80 and CD86 (Figure S22C).

### Atlas of sex differences in primary progressive MS peripheral blood mononuclear cells

We next explored the sex transcriptomic profiles of peripheral immune cells in PPMS (Figure 5A). Figure S24 reports marker gene expression, cell distribution by group and sex, and the number of statistically significant features, while Figures S25 to S29 summarize the results for each cell type. We identified significant functions in broad biological categories with similar proportions between males and females (Figure 5B); revealing sex alterations in the immune transcriptomic profile that impact several functional layers (Figure 5C). While the innate system suffered consistent proportional changes in NK cells and monocytes, we observed more notable differences for CD8+ T cells than CD4+ T cells in the adaptive immune system. We also encountered sex differences affecting the activation of signaling pathway effectors (Figure S30) and the strength of interaction between ligand-receptor pairs (Figure S31). In the latter, we noted a lack of commonly activated pathways in MS males (vs. control males) and MS females (vs. control females).

**Figure 5.**
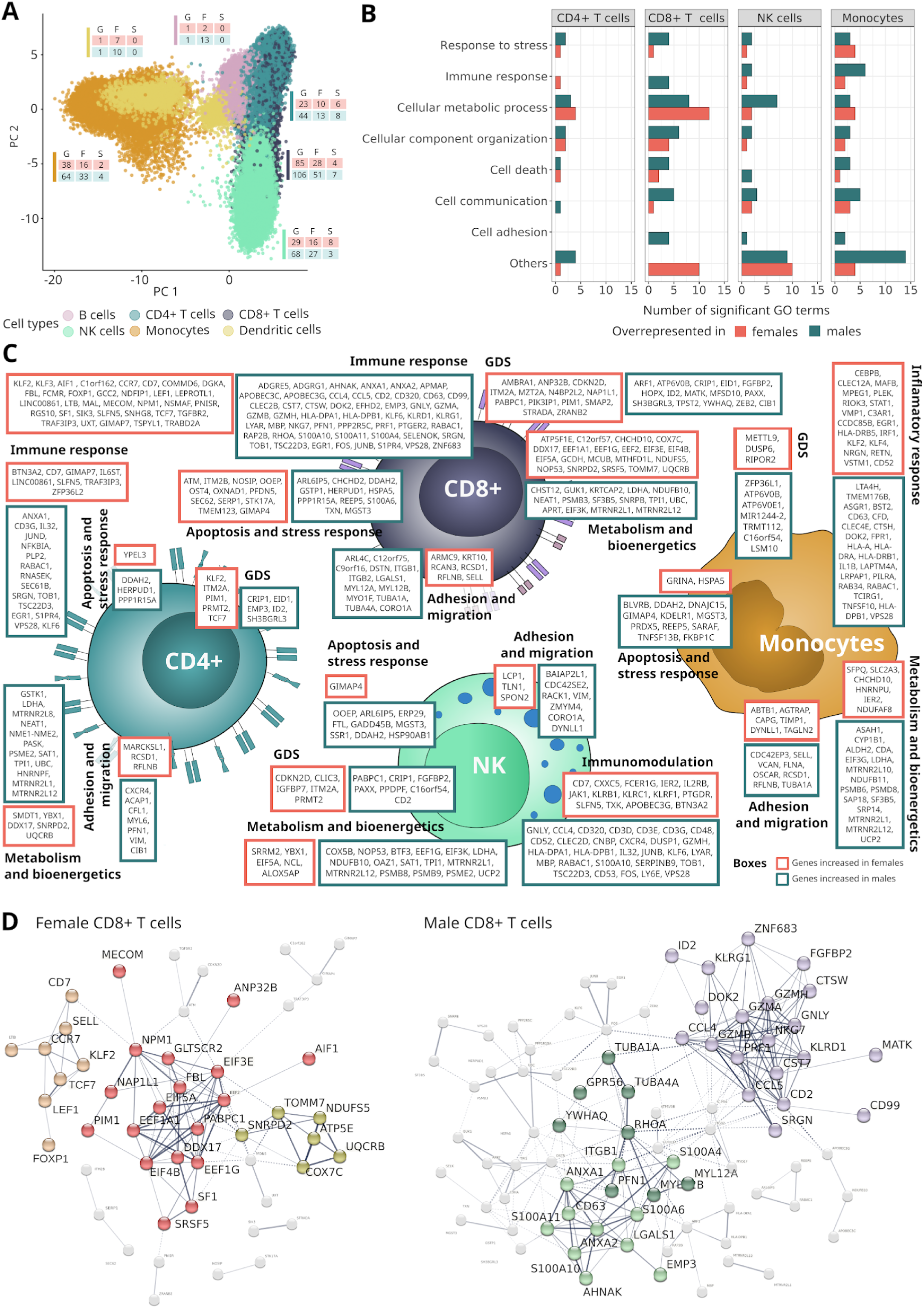
Transcriptomic landscape of sex differences in peripheral blood mononuclear cells of primary progressive MS patients. (**A**) Cell type distribution in PCA dimensions. Each dot represents a cell, colored by the annotated cell type. Colored boxes indicate that the number of genes (G), functions (F), and signaling pathways (S) significantly increased in females (orange) and males (blue). (**B**) The number of significant BP-GO terms by cell type classified under broader biological categories. (C) Atlas of sex differences in gene expression patterns. Genes classified in general biological terms. Colored boxes indicate the sex patterns: genes increased in females (orange) and males (green). D and E) Protein-protein interaction networks for (D) Female and (E) male CD8+ T cells. Edge thickness indicates the structural and functional confidence of the interaction, representing the intra-group (solid line) and inter-group (dotted line) connections. Clusters of interest are highlighted with colors. *BP-GO: Biological processes from Gene Ontology; MS: multiple sclerosis*.

#### Marked sex differences in CD8+ T cells of primary progressive MS describe predominant cytolysis in males and homeostasis processes in females

CD8+ T cells exhibited the most pronounced sex differences as previously described (Figure 5A-C). Genes with increased expression in females and males in CD8+ T cells formed significant interaction networks (p-values of 3.84e-14 and 1.0e-16, respectively) (Figure 5D). We infer that female CD8+ T cells could restore cellular homeostasis to a greater degree than males, as females displayed the increased expression of genes related to the regulation of protein translation (red cluster), the differentiation and survival processes of T lymphocytes (ochre cluster), and energy production through oxidative phosphorylation and mitochondrial maintenance (yellow cluster). Male CD8+ T cells may exhibit a more activated and harmful state thanks to higher cytolytic responses through granzyme and perforin-mediated apoptotic processes (purple cluster), calcium regulation (light green cluster), and the formation of extracellular vesicles (park green cluster).

### Sex differences in immune system status cluster relapsing-remitting MS and primary progressive MS cell types

After independently delineating sex differential patterns in immune cells for RRMS and PPMS, we sought to explore the significant disparities across these MS subtypes. To this end, we clustered the immune cells from both subtypes (CD4+ T cells, CD8+ T cells NK cells and monocytes) based on the expression profiles of the genes previously identified as significantly different between sexes. We performed the clustering analysis using the gene expression profiles with broader sex differential patterns, that is, genes significantly different at least in three of the eight cell types evaluated. This analysis revealed an initial stratification of cells primarily by disease subtype (RRMS or PPMS), followed by further stratification based on class (adaptive or innate immune system) (Figure 6). These findings suggest that critical disparities in the immune system impact the clinical variability observed between MS subtypes rather than subtle variations within a specific cell type. The 67 genes that supported clustering formed a highly connected interaction network (p-value: < 1.0e-16, Figure S32) primarily related to stimuli responses such as reactive oxygen species, cytokines, lipids, and leukocyte differentiation.

**Figure 6.**
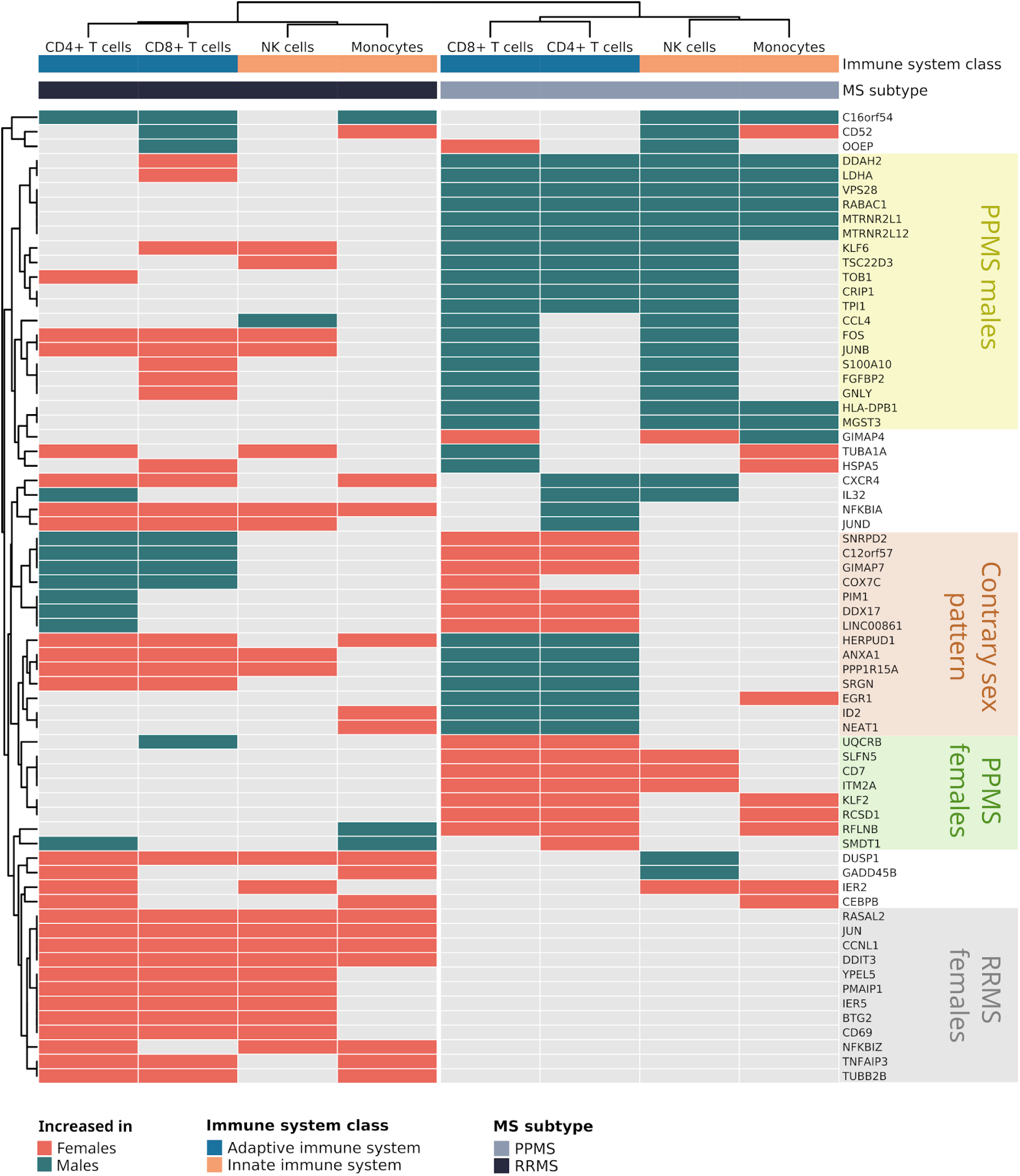
Clustering of relapsing-remitting MS and primary progressive MS immune system cell types based on their sex differential profile. Heatmap showing the classification of CD4+ T cells, CD8+ T cells, NK cells, and monocytes in RRMS and PPMS subtypes (columns) from genes with significant expression by sex in at least three of the evaluated cell types (rows). Disease subtypes, immune system classification (innate or adaptive), and the sex where the significant increment is detected (females or males) are specified. *NK: Natural killer; PPMS: primary progressive multiple sclerosis; RRMS: relapsing-remitting multiple sclerosis*.

RRMS significant genes displayed a more predominant increase in females, with a specific core that does not display significant dysregulation in PPMS (Figure 6, gray box). The RRMS female gene set reflects an activated state (CD69) driving responses, especially to stress and apoptosis (e.g., IER5, DDIT3, and BTG2). Conversely, the majority of significant PPMS genes are increased in males, with some exhibiting an increase in RRMS females (Figure 6, yellow box). A high proportion of PPMS genes increased in males relate to the modulation of inflammation in addition to processes not highlighted in the RRMS female core, such as glycolytic metabolism (LDHA and TPI1) and translation of mitochondrial proteins (MTRNR2L1 and MTRNR2L12). Interestingly, the adaptive immune system presented a gene set patterned inversely based on disease subtype, a scenario we did not observe in the innate immune system (Figure 6, orange box). Of note, ANXA1 and SRGN, which have relevance in CNS infiltration, suffer from increased expression in RRMS females and PPMS males.

### Sex differential viral responses and antigen presentation by MS subtype

Considering the extensive association of MS susceptibility with viral responses and specifically to genetic variants in the human leukocyte antigens (HLAs), we investigated the potential existence of sex differences in our data. RRMS females, compared to males, present more significant functions related to viral infection, while the opposite pattern occurred for PPMS. In detail, RRMS females exhibited the significant functions of *positive regulation by host of viral transcription* and *regulation of viral process* in CD4+ T cells, *cellular response to virus* in NK cells, and *negative regulation by host of viral transcription* in monocytes. We found no significant viral-related function in RRMS males. PPMS males displayed the significant functions of *positive regulation of defense response to virus by host* in CD8+ T cells, *response to virus* in NK cells, and *defense response to virus* in monocytes. The latter represents the only virus-related function overrepresented in PPMS females (also in monocytes). We observed the same tendency when evaluating the sex-differential expression pattern of significant genes belonging to these functions (Figure 7A).

**Figure 7.**
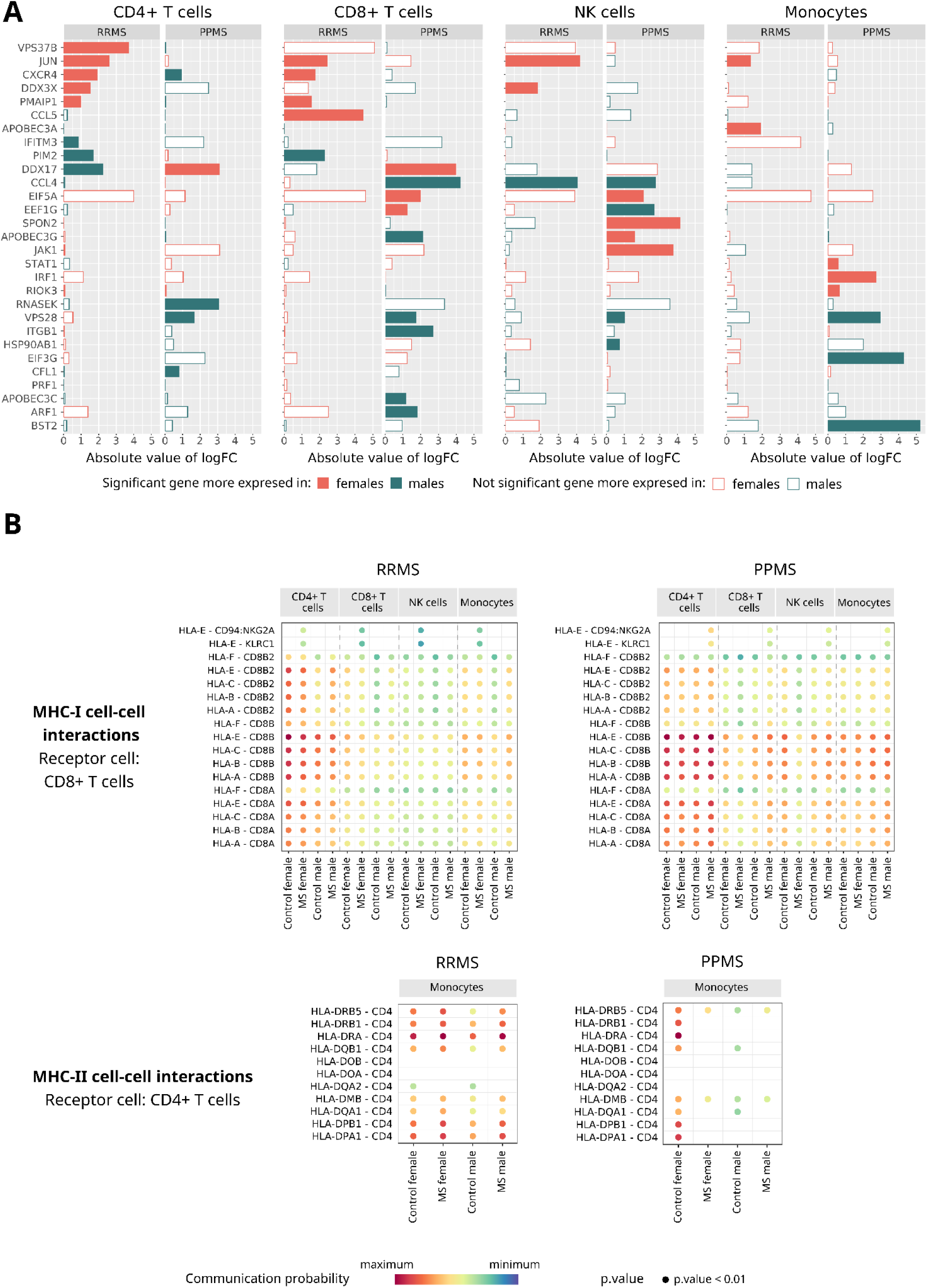
Sex differences in viral responses and antigen presentation between relapsing-remitting MS and primary progressive MS. (A) Patterns of gene expression alterations associated with viral responses by cell type and disease subtype. The magnitude of change in the SDID comparison (X-axis) for each gene extracted from the response to the virus and viral process (Y-axis) functions. Genes selected as significant in at least one cell type. (B) Significant ligand–receptor interactions involved in MHC-I (top) and MHC-II (bottom) signaling pathways, as inferred by CellChat analysis. Each dot represents a ligand–receptor pair with a statistically significant interaction. Dot color reflects the probability (i.e, strength) of the inferred interaction, with reddish colors representing higher probabilities. *MHC: Major histocompatibility complex; NK: natural killer; PPMS: primary-progressive multiple sclerosis; RRMS: relapsing-remitting multiple sclerosis; SDID: sex-differential impact of disease*.

HLA genes play a critical role in the immune response by presenting foreign antigens to T cells; furthermore, they influence MS susceptibility/protection by regulating the immune system’s response to self-antigens. We observed the notably increased expression of HLA-DRB5 in RRMS male monocytes and PPMS female monocytes (Table S6). While HLA-DRB5 represents the only HLA family gene with increased expression in PPMS females, we did observe increased expression of HLA-DR and HLA-DP genes in PPMS males (Table S6).

Major histocompatibility complexes (MHCs) are the protein products of HLA genes. Here, we identified sex differences in their communication strengths (Figure 7B). We encountered the major sex difference in MHC-I for RRMS and PPMS between the HLA-E ligand with the CD94:NKG2A and KLRC1 receptors (Figure 7B, top). These interactions became significant for RRMS females and PPMS males. We observed much stronger interactions for MHC-II for female controls than male controls at the baseline (Figure 7B bottom). RRMS males displayed increased interaction strengths in MHC-II to reach the same level as RRMS females, with both sexes demonstrating significant interactions for almost all ligand-receptor pairs (Figure 7B, bottom). Conversely, ligand-receptor interactions disappear in PPMS females compared to control females, while in PPMS males maintain a state similar to control males (Figure 7B, bottom).

### Atlas-MS web platform

To encourage a more comprehensive coverage of this transcriptomic landscape, we developed the interactive web tool https://bioinfo.cipf.es/cbl-atlas-ms/. This application offers a user-friendly interface that supports the free access and visualization of all the research outcomes, by which users can interactively explore the results by selecting the MS subtype, cell types, and molecular features of interest (Figure 8).

**Figure 8.**
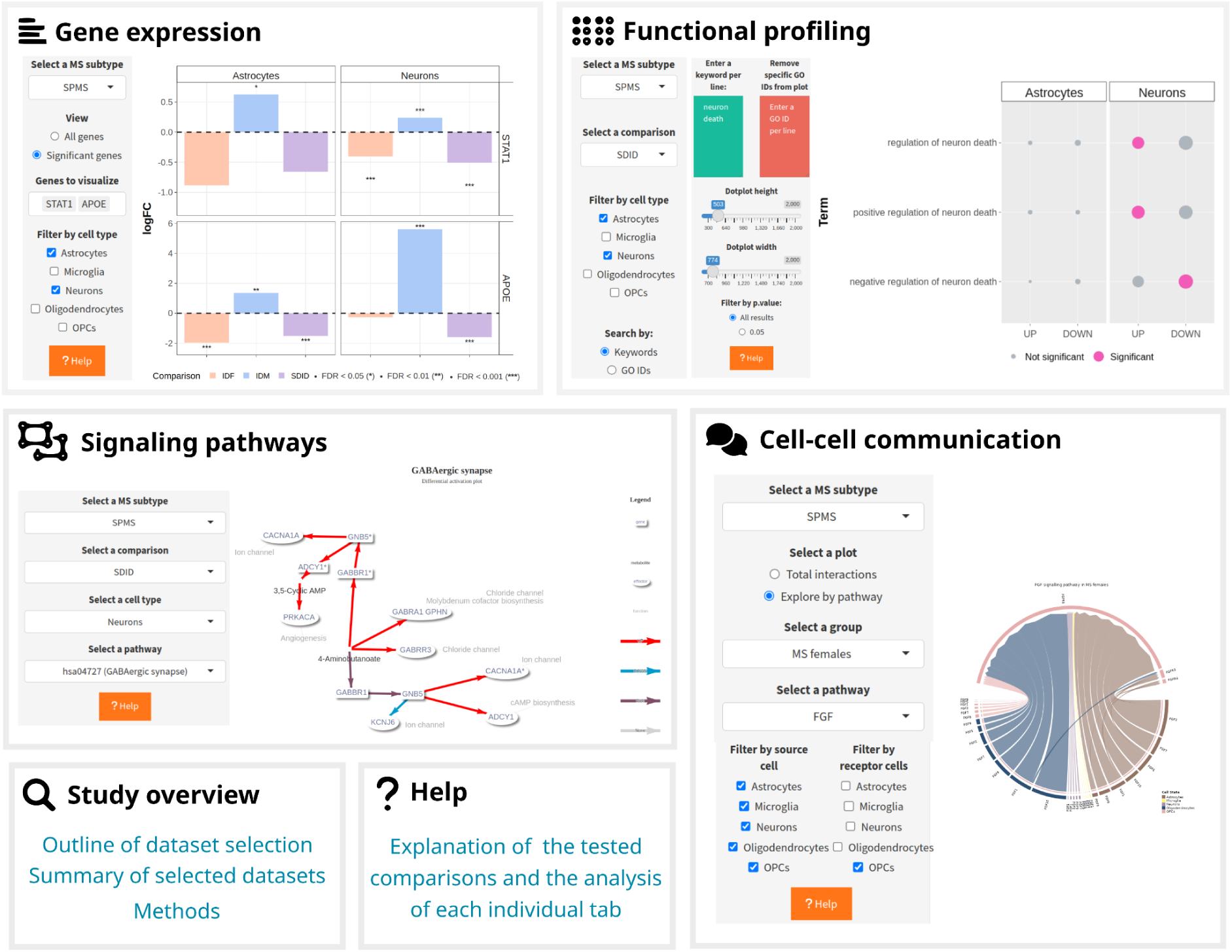
Atlas-MS web platform. The interface contains six tabs: *Differential Gene Expression* (to explore how gene expression change in MS based on being male, female and the underlying sex differences), *Functional profiling* (to delve into the biological functions in which dysregulated genes are involved), Signaling pathways (to visualize differential activation of pathways in MS based on being male, female and the underlying sex differences), *Cell-cell communication* (to examine cell-cell communication networks evidenced by condition and sex) *Study Overview* (to review the study design), and *Help* (to access user guidance)

## DISCUSSION

Despite the well-documented sex differences in MS progression, the specific underlying molecular mechanisms and the impact of determinant factors, such as sex, remain underexplored. This study outlines the sex differential single-cell transcriptomic landscape in MS subtypes. We hope to potentiate biomarker discovery and the development of personalized therapeutic strategies by providing a better comprehension of sex differential molecular mechanisms within each cell type. Representative findings were selected based on their biological relevance to each cell type, prioritizing contextually meaningful processes in the disease.

We first uncovered molecular mechanisms that may explain differences in CNS samples from SPMS. Excitotoxicity, that is, neuronal damage due to exposure to elevated glutamate levels, represents a major MS hallmark.^13^ We suggest that female neurons increased expression of genes that may act to compensate for such excitotoxicity, such as glutamate and GABA receptors and cation transporters to modulate excitability, and increased activation of GABA signaling effectors. We validated this pattern in an independent cohort, although further complementary analyses will be required for definitive confirmation. In contrast, male neurons appeared more susceptible thanks to the overexpression of genes encoding ATPases subunits that provide energy for vesicle biogenesis while exhibiting a higher activation of glutamate signaling effectors.

The glial-driven SPMS stress response represents a complex set of processes to different stimuli that account for sex differences; however, studies currently differ in describing their sex differential impact.^11^ We report that male astrocytes express increased levels of glutamate reuptake-related genes, which could modulate excitotoxicity in the synaptic cleft.^14^ Conversely, female astrocytes present key genes (such as HIF3A and EGLN3) in addressing the hypoxic stress characterizing MS demyelinating lesions.^15^ Female astrocytes also appear to balance their profile towards neuroprotection or reactivity in SPMS by regulating the expression of multifaceted genes via lncRNAs such as NEAT1 - a negative regulator of neuronal excitability in neurodegenerative diseases^16^ - and LINC-PINT - a potential regulator of oxidative stress in Parkinson’s disease substantia nigra.^17^

Previous studies have demonstrated more effective remyelination in aged female rodents than males^18^ and a hormonal sex-dependent regulation of myelin repair markers in experimental autoimmune encephalomyelitis.^19^ Expanding on these findings, our research revealed sex differences in myelin clearance and repair in SPMS, which could be crucial to developing effective remyelination therapy strategies. Female oligodendrocytes expressed a greater number of neuronal-like genes than males, which would be relevant in establishing axon-myelin contacts.^20^ Female OPCs also increased the expression of QKI, a pivotal gene that promotes oligodendrocyte differentiation^21^ and maintains myelin sheaths by regulating lipid homeostasis.^22^ To cope with myelin damage, PDGF and FGF signaling represent crucial signals. PDGF enhances the growth, differentiation, and survival of OPCs.^23^ The interaction between PDGFA (OPCs) - PDGFRA (OPCs) emerged in SPMS males when compared to male controls (Figure 3A), which represents a crucial interaction when considering recovery from chronic demyelination.^24^ While this interaction becomes established in both control females and MS females, the PDGFC (OPCs) - PDGFRA (OPCs) interaction intensifies in SPMS females, which relates to reduced neuroinflammation in MS.^25^ For FGF signaling, which promotes myelination, SPMS females suffered from an increase in the proportion of OPC-neuron interactions that could support myelin sheath establishment. Further studies will be necessary to determine whether these transcriptional changes translate into measurable effects on OPC maturation and remyelination capacity. We also found that SPMS female microglia displayed more genes related to phagocytosis and lipid metabolism, which could favor the clearance of myelin.^26^ Meanwhile, male microglia presented increased SCARB1 gene expression, which clears amyloid-β deposition in Alzheimer’s disease.^27^ Of note, male astrocytes presented increased levels of APOE4, which potentially promotes lipid droplet accumulation, compromising their functionality to offer metabolic support to neurons.^28^ Interestingly, APOE4 is associated with an increased tendency of MS progression^29^ and cognitive impairment.^30^

Previous efforts have been made to understand the role and status of the peripheral immune system. The Multiple Sclerosis Competence Network underscored the importance of investigating the MS peripheral immune system by reporting three different blood endophenotypes linked to distinct disease outcomes.^31^ However, the evidence related to sex differential disturbances remains inconclusive. We reported a greater number of significant features in RRMS females than males (Figure 4). These features participate in critical processes such as inflammation, cell signaling, and cell-cell adhesion, which may prompt exacerbated immune responses and induce higher female susceptibility to MS. We interestingly identified the genes encoding the AP-1 subunits (FOS, JUN, JUNB, and JUND) forming the central hub of the female immune network (Figure 4E), reflecting their importance as master regulators of a broad range of immune system processes such as proliferation, CNS infiltration, and inflammation.^32^ This set of genes could serve as a starting point to explore immune targets differentially regulated between sexes. We also identified the NFKBIA and NFKBIZ genes (encoding inhibitors of the MS-associated NF-κB - a pathway^33^ that may counteract the proinflammatory status). Interestingly, these findings are likely driven by sexual hormones, as estrogen promotes AP-1 complex activation^34^ and NFKBIA gene and protein expression.^35^

Our findings also suggest that the adaptive immune system of RRMS males suffers from the enhanced activity of the mitochondrial electron transport chain (Figure 4C). Mitochondrial DNA variants have also been proposed as susceptibility biomarkers of RRMS.^36^ Regarding PBMCs, RRMS patients exhibited mitochondrial dysfunction compared to controls, which becomes more pronounced in patients experiencing a more aggressive disease course.^37^ These findings led us to hypothesize that exploring the mitochondrial status of CNS-infiltrating peripheral T lymphocytes may provide insight into the more rapid neurodegeneration observed in MS males.

The role of the peripheral immune system in PPMS remains less well documented. PPMS differs from other subtypes; treatments effective in RRMS have reduced efficacy in PPMS^38^, and the expression of clinical MS biomarkers remains lower in PPMS compared to SPMS.^39^ These findings suggest the necessity to comprehensively characterize the PPMS immune profile. In contrast to RRMS, where alterations in females predominate, we found that PPMS males and females have a similar proportion of enriched functions involving broad immunophenotypic features (Figure 5B). Interestingly, a higher number of functional and gene expression differences according to sex occur in CD8+ T cells compared to other cell types. Intrathecal CD8+ T cell subtypes represent critical elements in the immunopathogenesis of PPMS and associate with white matter injury and thalamic atrophy.^40^ They hold particular interest in PPMS due to the significant decline in peripheral CD8+ T cell number observed with age in affected patients compared to healthy controls, suggesting an accelerated aging effect in MS.^41^ We reported sex-biased CD8+ T cell responses (Figure 5D), with PPMS females more likely to restore cellular homeostasis than males and PPMS males exhibiting a more activated state characterized by heightened cytolytic responses. These responses may represent an explanation for the more severe neurodegeneration observed in MS males, triggering more rapid progression.

We identified a genetic signature composed of 67 genes that, based on their sex differential profile, classified the MS immune system by disease subtype (Figure 6). This signature may enable the characterization of progression-related biomarkers and the development of tailored treatment approaches by identifying specific subtypes in MS patients. 12 genes that undergo significantly increased expression in female RRMS but remain not significant in PPMS mainly relate to modulating inflammation, a key differentiating factor between RRMS (higher inflammation) and PPMS (lower inflammation). Of them, CD69 protein has been detected in MS lesions in both brain and infiltrated immune cell types.^42,43^ Therefore, increased CD69 expression in RRMS females holds promise for further investigation. Meanwhile, PPMS-associated genes primarily increase in male patients. Although many also play roles involved in inflammation, glycolytic metabolism stands out, particularly through the expression of the TPI1 (catalyzes the isomerization of glyceraldehyde 3-phosphate and dihydroxy-acetone phosphate) and LDHA (catalyzes the reversible conversion of pyruvate to lactate) genes. Peripheral immune system cells exhibit metabolic dysfunction, partly due to their activation during immune responses and the mitochondrial damage they undergo.^44^ This immune-metabolic phenomenon remains a current field of interest in exploring therapeutic targets. Given our findings, we emphasize the importance of elucidating sex-specific metabolic profiles, which may offer valuable insights for targeted therapeutic interventions.

The strongest MS genetic associations locate to the HLA gene loci, which encode proteins crucial for the immune system’s ability to distinguish between foreign organisms and our cells.^45^ This gene family has been implicated in several autoimmune conditions.^46^ Furthermore, distinct sex-specific differences in HLA expression have been documented across various contexts, including peripheral immune responses to lipopolysaccharide^47^ and generalized aggressive periodontitis.^48^ MS is no exception, as we observed significant variations in HLA gene expression levels and ligand-receptor interaction strengths within their antigen presentation roles (Figure 7). We highlight expression changes in the HLA-DRB5 gene, which increases in RRMS females and PPMS males. Demyelinated lesions are characterized by increased expression of HLA-DR genes^49^; HLA-DRB5 plays an intricate role in MS as it may mitigate disease severity while actively presenting antigens derived from myelin promoting autoimmunity.^50,51^ Additionally, we found that HLA-E protein interactions with CD8+ T cells became significant in RRMS females and PPMS males, which may play a pivotal role in presenting autoantigens.^52^ If such interactions differ between subtypes, ultimately they may contribute to disease progression, potentially exacerbating the relapses in females (RRMS - subtype characterized by inflammation) and the rapid neurodegeneration in males (PPMS - subtype characterized by less inflammation but continuous neurodegeneration). Regarding HLA class II genes, which also influence genetic susceptibility to MS^53^, RRMS males exhibited heightened interaction strengths, reaching levels observed in RRMS females. In contrast, ligand-receptor interactions disappeared in PPMS females compared to female controls, while PPMS males maintained similar patterns to control males. This loss of intensity in connections in female PPMS could represent an area of further exploration to elucidate the reason for higher RRMS but not PPMS susceptibility in females. Moreover, it may point to a potential mechanism allowing that, although females have a more active immune system in healthy conditions, this is attenuated to have a neurodegenerative centered progression in the PPMS subtype.

We acknowledge the strengths and limitations associated with this study. We performed an in-silico approach underscoring the importance of promoting Open Science initiatives and adhering to FAIR (Findable, Accessible, Interoperable, Reusable) principles to encourage data reuse and enhance the advancement of research. Although this strategy captured the state-of-the-art scRNA-seq MS data in humans, the heterogeneous characteristics among the retrieved datasets hindered analysis. We wish for additional data to analyze novel combinations of tissue and MS subtypes. Additional data will also improve the robustness of our current findings, which are limited by the low number of cells or imbalance among groups in some populations like microglia, OPCs, and oligodendrocytes. Moreover, it would enable the inclusion of relevant cell types, such as B cells and dendritic cells, that were excluded due to insufficient representation. Along with differential expression and functional profiling analyses, we inferred signaling pathway activation and cell–cell communication from transcriptomic data. These results offer additional insight into the molecular landscapes, although confirmation using complementary approaches would be valuable. Since these analyses rely on predefined pathway databases, future studies could explore non-canonical signaling differences.

We previously characterized sex differences in MS by analyzing microarray and bulk RNA-seq transcriptomic data in brain and blood samples.^54^ This research identified sex differential gene expression levels averaged across all cells within a sample; however, our current work can dissect cellular heterogeneity. Thus, we shed light on cell type-specific responses. The continued dissection of differential expression profiles in additional subpopulations remains of great interest to refine our understanding of sex differential molecular mechanisms. Although we attempted to cover diverse areas of disease pathology in the CNS and immune system, addressing all encountered differences within a single manuscript remains unfeasible. To provide broader and free access to our findings, we developed a user-friendly website based on Shiny (https://bioinfo.cipf.es/cbl-atlas-ms/), enabling any interested user’s interactive exploration of the complete results.

Overall, we hope to have contributed to the characterization of MS molecular pathology at the single-cell level by facilitating the search for potential biomarkers and the development of novel targeted therapies while considering the MS subtype and the individual’s sex. To the best of our knowledge, we generated the first single-cell sex differential transcriptomic atlas of MS, providing results by tissue and disease subtype. Our investigation identified genes, functions, and pathways worthy of consideration in the potential development of biomarkers and targeted therapies.

## CONCLUSIONS

We have identified sex differences in the central nervous system and peripheral immune cell types in the spectrum of multiple sclerosis. The higher relapse frequency and incidence in females during early stages (RRMS) may be driven by an increased expression of multifaceted genes that would exacerbate inflammatory responses. As the disease progresses, males experience faster CNS neurodegeneration. We propose it may be linked to compensatory mechanisms present in SPMS females, which would help to cope with excitotoxicity (in neurons) and to promote myelin clearance (in glial cells), together with the increased cytotoxic activity of male PPMS CD8+ T cells. The sex-differential profile of the immune cells also allowed us to classify cell types based on their clinical subtype, obtaining a transcriptomic signature that could help to determine the status of multiple sclerosis patients. Overall, we hope our findings could act as a valuable resource to promote further research in understanding multiple sclerosis considering the sex of the individuals.

## METHODS

### Study design and resource availability

Literature screening was conducted in September 2025 following a systematic search criteria for dataset identification. Inclusion criteria were implemented using the keywords “multiple sclerosis,” “Homo sapiens,” “single cell,” or “single nuclei” or “single nucleus” in the GEO, ArrayExpress and UCSC Cell Browser public databases. Web browser searching was also performed.

Of the datasets identified, three were analyzed to generate the sex-differential landscape, while the grey matter neurons annotated in the study by Macnair *et al*.^12^ were used for validation of CNS results. The processing of this dataset was the same as for the others, except that we retained the original authors’ annotation.

The identified studies were manually filtered using the following exclusion criteria: i) data type: not containing scRNA-seq or snRNA-seq data, ii) disease examined: studies not based on MS, iii) experimental design: required MS patients as cases and healthy individuals as controls, iv) sex: sex as a variable not registered or not available, v) sample count: at least data from three different individuals at each condition and sex (control females, MS females, control males, MS males), and vi) unavailability of the gene expression matrix or metadata files. Finally, raw count matrices and sample metadata were downloaded for each selected study.

Bioinformatics analyses were individually performed for each selected study. Firstly, the data were processed as follows: i) quality control filtering, ii) normalization, iii) highly variable gene (HVG) selection, iv) dimensionality reduction, v) cell identity clustering, vi) cell type annotation, and vii) gene marker evaluation. Results were then obtained for each cell type in females, males, and MS sex differences by exploring statistical changes in gene expression patterns, functional profiles, signaling pathways, and cell-cell communication.

The bioinformatics code was developed using R (version 4.1.2). Table S7 provides a record of the versions of corresponding R packages. Bioinformatic code is available at https://github.com/IrSoler/cbl-atlas-ms.

### Data acquisition, preprocessing, and cell-type annotation

Gene expression matrix and metadata files were downloaded to develop individual analyses. Standardization of the gene nomenclature was ensured, verifying that Gene Symbols were available in all datasets. Next, a quality control analysis was performed to incorporate data from viable cells/nuclei in downstream processing. This step was adapted to suit the singularities of each dataset, maintaining homogeneity among studies as much as possible. The following parameters were calculated by sample: number of cells/nuclei, library size, number of expressed genes, and mitochondrial gene ratio. The graphical distribution of each parameter was represented before and after filtering to explore the need to eliminate samples. Pertinent cutoffs were implemented to filter poor-quality cells/nuclei. For the UCSC-MS dataset, nuclei with less than 1000 library size, less than 500 genes expressed, anomalous values (outliers) for mitochondrial gene ratio identified with isOutlier function from scuttle R package^56^, or nuclei aggregates detected with scDblFinder R package (https://github.com/plger/scDblFinder) were removed. Quality control analysis was also performed at the gene level, discarding genes expressed in less than three nuclei or genes other than protein-coding/micro/long non-coding RNAs. HUGO Gene Nomenclature Committee guidelines were followed to identify these gene types. The graphical distribution of the parameters before filtering revealed that GSE144744 cohorts were previously filtered by the authors, corroborating the absence of outliers; therefore, additional filtering steps were not performed.

Filtered count matrices were normalized using a deconvolution approach^57^ and transformed into a log2 scale. Each gene’s biological and technical variability was estimated, considering the batch effects reported in the metadata files. 20% of HVGs based on biological heterogeneity were selected. All steps were executed using the scran R package.^58^ HVG expression was used as input data to implement dimensional reduction approaches that further compact expression profiles. In detail, summarization was assessed by PCA (principal component analysis) using the scater R package.^56^ The relevant principal components were selected by applying the elbow point method with PCAtools R package (https://github.com/kevinblighe/PCAtools). Moreover, the non-linear dimensionality reduction strategies tSNE (T-distributed stochastic neighbor embedding) and UMAP (uniform manifold approximation and projection) were computed for visualization.

Cells were classified based on the similarity of the previously selected principal components. To this end, a graph built on the Shared Nearest Neighbours algorithm was constructed using the scran R package.^58^ Groups were defined with the walktrap algorithm (*k* = 20) with the igraph R package.^59^ Cell type was assigned to each cell/nucleus by comparing expression profiles to public references. The BRETIGEA R package^68^ was used for nervous tissue (UCSC-MS dataset), and the SingleR package^60^ for peripheral blood mononuclear cells (GSE144744 cohorts). The MonacoImmuneData reference was selected from the celldex R package^60^ as a standard reference for analyzing human peripheral blood mononuclear samples. Unassigned cells or cells not classified as major cell types from the CNS (neurons, astrocytes, microglia, oligodendrocytes, OPCs) and blood (CD4+ T cells, CD8+ T cells, NK cells, monocytes, B cells, dendritic cells), respectively, were discarded. For the CNS dataset, we next confirmed the absence of expression of marker genes for infiltrated immune cells: CD169 (macrophages), CD3 (T cells), CD19 and CD20 (B cells). As an additional quality control measure, following annotation with reference datasets, we performed differential expression analyses of marker genes: astrocytes (GJA1, PRDM16, SLCO1C1, ALDH1L1, EYA1), microglia (C3, CD74, C1QB, KCNK13, GPR34), neurons (GABR2, HCN1, LPHN2, KCNJ3, PCSK2), oligodendrocytes (OPALIN, MAG, MOG, ERMN, ANLN), OPCs (COL9A1, GPR17, PDGFRA, STK32A, CALCRL), B cells (CD22, CD79A, CD79B, MS4A1, BANK1), CD4+ T cells (IL7R, MAL, IL6ST, TCF7, AQP3), CD8+ T cells (CD8A, CD8B, CD8B2, CD3D, CD3G), NK cells (NKGG7, CTSW, GZMA, FCGR3A, KLRD1), dendritic cells (CLEC10A, IRF7, CCDC88A, FCER1A, ITM2C), and monocytes (CD14, CD36, S100A9, S100A8, MS4A6A). Gene expression distributions can be consulted in Supplementary Figures S5-A, S16-A, and S24-A. Then, genes overexpressed in each cell type compared to the remaining cells were screened by implementing the statistical Wilcoxon test with the scran R package.^58^ Adjusted p-values were calculated by the Benjamini-Hochberg method^69^, considering genes significant when attaining a false discovery rate (FDR) < 0.05. Significantly overexpressed genes by cell type were corroborated as marker genes described in scientific literature, increasing the robustness of the annotation.

We also accounted for potential sources of variability. Supplementary Figures S2–S4G illustrate that the primary contributor to variation in the first principal components is cell type assignment. Supplementary Figure S33 presents the distribution of cells per sample (donor), ensuring that all cell types are represented across all individuals. Covariates included in the differential expression model are described in the section *Differential gene expression analysis*.

### Evaluated comparisons

Differential gene expression and functional analyses were performed to statistically identify differences in MS from a sex perspective (see following sections), testing three independent comparisons for each cell type analyzed. The MS.Female - Control.Female comparison assessed the impact of MS in females (IDF), revealing the differences between MS females and healthy individuals. The MS.Male - Control.Male comparison reported the impact of MS in males (IDM), revealing the differences between MS males and healthy individuals. The (MS.Female - Control.Female) - (MS.Male - Control.Male) comparison evaluated the sex differential impact of disease (SDID), which aimed to identify sex differences among MS patients while considering inherent sex variability in healthy individuals.

Significant characteristics were selected in all three comparisons to explore the biological results.

### Differential gene expression analysis

A hurdle model was implemented using the MAST R package.^61^ A logistic regression was considered to model if a gene would be expressed (discrete distribution), and a conditioned Gaussian linear model to represent the expression level (continuous distribution).

For each analysis, the model included a variable representing the condition and sex conforming to four groups (control females, MS females, control males, MS males) and another variable containing the scaled number of expressed genes (MAST recommendation). Additional covariables were added to control their variability minimizing the effect of confounders in gene expression patterns. UCSC-MS comparisons incorporated sample, age, affected cerebral region, capture batch effect, sequencing batch effect, and cell cycle state. For the GSE144744 cohorts, the variables sample, age, previous treatments, batch effect, and cell cycle state were reported. The cell cycle state represents a source of variability not provided in corresponding metadata files in any dataset, so it was calculated using the scran R package.^62^

A likelihood-ratio test was then performed to calculate each dataset’s statistical parameters by cell type and comparison.

### Functional profiling

#### Inference of protein-protein interaction networks

Protein-protein interaction analyses were performed using the STRINGdb R package.^63^ Interaction networks were obtained by keeping default parameters and examining the database’s total number of physical and functional interactions. For visualization purposes, weighted edges were represented based on the confidence of the interaction (the more significant the thickness, the greater the confidence), hiding disconnected proteins.

#### Functional overrepresentation analysis

Altered biological processes from Gene Ontology (BP-GO) were identified through a modular enrichment analysis implemented using the weight01 algorithm from the topGO R package.^64^ Significant genes were evaluated for the BP-GO terms obtained from the org.Hs.eg.db R package. Significant terms were clustered, and a word cloud was generated for each cluster. This process was implemented with the simplifyEnrichment and simplifyGO R packages.^65^ Semantic similarity matrices were obtained by grouping terms using the Louvain clustering method. Word clouds were created, with the size of each word proportional to frequency. Visual atlases were created of significantly dysregulated genes in each cell type. Genes were grouped into general functional categories based on BP-GO terms and associated functional descriptions from the literature.

### Signaling pathways analysis

Statistical differences in signaling pathway activity were identified using the hipathia R package.^66^ Signal transduction was calculated for each effector subpathway of each pathway from the Kyoto Encyclopedia of Genes and Genomes (KEGG).^67^ Differential effector activation analysis was performed using the MAST R package^61^, following the specifications described in the differential gene expression analysis. Suitable pathways for analysis were selected. Criteria were established based on the type of tissue sample (Table S8).

### Cell-cell communication analysis

The R package CellChat^68^ inferred interactions between ligand-receptor pairs. The CellChatDB human database screened 1939 ligand-receptor interaction pairs involving 546 ligands and 507 receptors. Communication probabilities were computed for each interaction between pairs of cell types in each group of interest (control females, MS females, control males, and MS males), with these values measuring the interaction strength. The direction of interactions was assessed by establishing the cell type that provides the ligand and/or the receptor. Cellular communication networks were then calculated by counting the total number of interactions and the differential number of interactions among females (MS females vs. control females comparison) and males (MS males vs. control males comparison).

### Web platform development

Our web tool (https://bioinfo.cipf.es/cbl-atlas-ms/) was developed using the structure of the R shiny package (https://github.com/rstudio/shiny, https://shiny.posit.co/).

### Statistical analysis

Corresponding statistics to each analysis were determined. Adjusted p-values were calculated by the Benjamini-Hochberg method^69^. LogFC values were determined to reveal the magnitude of the differential expression (logFC absolute value) and in which direction the genes were more significantly expressed (logFC sign). For guidance on interpreting the logFC, refer to our website’s “Help” section: https://bioinfo.cipf.es/cbl-atlas-ms/. Features were considered significant when attaining a false discovery rate < 0.05, adding the cut-off logFC absolute value > 0.5 to differential gene expression results.

## ABBREVIATIONS

BP-GO: biological processes from the Gene Ontology
CNS: central nervous system
IDF: impact of disease in females
IDM: impact of disease in males
FDR: false discovery rate
FGF: Fibroblast growth factor
GABA: gamma-aminobutyric acid GEO Gene Expression Omnibus HLA human leukocyte antigen
HVG: high variable genes
KEGG: Kyoto Encyclopedia of Genes and Genomes
logFC: logarithm of fold change
lncRNA: long non coding RNA
MHC: major histocompatibility complex
MS: multiple sclerosis
OPCs: oligodendrocyte precursor cells
PCA: principal component analysis
PDGF: Platelet-derived Growth Factor
PPI: protein-protein interaction network
PPMS: primary progressive multiple sclerosis
RRMS: relapsing-remitting multiple sclerosis
scRNA-seq: single-cell RNA sequencing
snRNA-seq: single-nucleus RNA sequencing
SDID: sex-differential impact of disease
SPMS: secondary progressive multiple sclerosis
tSNE: T-distributed stochastic neighbor embedding
UMAP: uniform manifold approximation and projection

## DECLARATIONS

### Ethics approval and consent to participate

Not applicable.

### Consent for publication

Not applicable.

### Availability of data and materials

The datasets analyzed in this work are publicly available: the SPMS postmortem brain tissue dataset was retrieved from Schirmer *et al*^69^, and the RRMS and PPMS peripheral immune cells datasets from Kaufmann *et al*^70^. The bioinformatic code can be found at https://github.com/IrSoler/cbl-atlas-ms and the complete results can be explored at https://bioinfo.cipf.es/cbl-atlas-ms/.

### Author contributions

Conceptualization: ISS, MRH, FGG; Data Curation: ISS; Investigation: ISS, BGC, RGR, CGR, LBM, HC, MIV, SGP, VT, MRH, FGG; Bioinformatic Analysis: ISS, BGC, RGR; Supervision: MRH, FGG; Writing-Original Draft Preparation: ISS, BGC, RGR, CGR, LBM, HC, MIV, SGP, VT, MRH, FGG. All authors read and approved the final manuscript.

## Acknowledgements

The authors thank the Principe Felipe Research Center (CIPF) for providing access to the cluster, co-funded by European Regional Development Funds (FEDER) to the Valencian Community 2014-2020. The authors also thank Stuart P. Atkinson for reviewing the manuscript.

## Supplementary Information

Supplementary information includes Supplementary Note, Tables S1-S8 for multiple supplementary tables and Figure S1-S32 for multiple supplementary figures.

